# Lethal Disruption of the Symbiotic Gut Community in Eastern Subterranean Termite Caused by Boric Acid

**DOI:** 10.1101/2024.06.26.600876

**Authors:** Aaron R. Ashbrook, Melbert Schwarz, Coby Schal, Aram Mikaelyan

## Abstract

The Eastern subterranean termite, *Reticulitermes flavipes* (Kollar), is a significant pest, causing extensive damage to structures that amount to substantial economic losses. Traditional termite control methods have utilized boric acid, known for its broad-spectrum insecticidal properties, yet its impact on termite gut microbiomes and the implications of such effects remain understudied. Our study evaluates the dose-dependent mortality of *R. flavipes* upon being provided boric acid treated filter papers and investigates the resulting dysbiosis within the termite gut microbiome. Consistent with reports from other insects, mortality increased in a dose-dependent manner, with the highest boric acid concentration (203.7 µg/cm^2^ of filter paper) significantly reducing termite survival. 16S rRNA gene sequencing of the gut microbiome revealed notable shifts in composition, indicating boric acid-induced dysbiosis. Aside from an overall decrease in microbial diversity, the relative abundance of some symbionts essential for termite nutrition decreased in response to higher boric acid concentrations, while several putative pathogens increased. Our findings extend the understanding of boric acid’s mode of action in termites, emphasizing its effect beyond direct toxicity to include significant microbiome modulation that can have dire effects on termite biology. Considering its potential to induce dysbiosis and potentially augment the effectiveness of entomopathogens, our study supports the continued use of boric acid and related compounds for termite-resistant treatments for wood.

## Introduction

The Eastern subterranean termite, *Reticulitermes flavipes* (Kollar) (Blattodea: Rhinotermitidae), is a major wood-destroying pest with a wide distribution in the U.S. and around the world (Dedeine et al. 2016). Subterranean termites display cryptic nesting behaviors, which often allow their colonies to go undetected when causing damage to structural wood (Oi 2022). Globally, termites impose an astonishing economic burden, with damages estimated at approximately USD 40 billion annually, with 80% of the damage caused by subterranean species (Rust and Su 2012; Oi 2022). The ability of *R. flavipes* and other termite species to inflict significant damage on wooden structures is attributed largely to their intricate symbiotic associations with gut microbiome (Brune 2014), which include protozoa (Gile 2023), bacteria (Brune and Dietrich 2015), and archaea (Protasov et al. 2023). This complex digestive symbiosis enables termites to effectively break down lignocellulose (Watanabe and Tokuda 2010), critically weakening the integrity of wood, and ultimately making structures more susceptible to collapse (Scharf 2020).

The gut microbiome of termites is a complex and dynamic system that is susceptible to disruption by antibiotics, as has been observed in *Zootermopsis nevadensis* Hagen (Blattodea: Archotermopsidae) and *R. flavipes* (Rosengaus et al. 2011). Force-feeding starch to *R. flavipes* can precipitate a dramatic shift in microbiome composition and functionality, chiefly characterized by the elimination of cellulolytic protozoa and their associated bacteria (Ikeda-Ohtsubo et al. 2010; Mikaelyan et al. 2017). This vulnerability creates the potential for utilizing disruptive agents that could even enhance the efficacy of pesticides, including biological control agents, by targeting and altering the termite gut microbiome.

Boric acid is an inexpensive, broad-spectrum insecticide that has been widely used to protect wood from pest damage and can cause mortality in termites (Gentz and Grace 2006; Schubert 2015). While the full scope and mode of its action remain to be fully elucidated, it has been demonstrated to negatively impact digestion and nutrient absorption in a broad range of insects (Cochran 1995; Klotz et al. 2002; Habes et al. 2006; Gwokyalya and Altuntaş 2019). Notably in termites such as *R. flavipes* and *Coptotermes formosanus* Shiraki (Blattodea: Rhinotermitidae), preliminary studies have demonstrated that boric acid eliminates gut protozoa (Kard 2001), impairing termite digestion, and reinforcing its reputation as a “stomach poison” (Ebeling 1978). However, the specific changes in the termite gut microbiome induced by boric acid remain under-explored.

Recent research on cockroaches, which share a close phylogenetic relationship with termites and have similarly complex gut microbiomes, highlights the disruptive effects of boric acid on these communities (Jiang et al. 2021; Yang et al. 2021). Inspired by these insights, our study seeks to examine how boric acid alters the gut microbiome composition of *R. flavipes*, particularly in relation to its lethal and sublethal impacts. We aim to bridge this gap by focusing on two objectives: 1) Characterizing the mortality of *R. flavipes* in response to various boric acid concentrations through feeding experiments, and 2) Investigating whether boric acid-induced disruption increases the prevalence of opportunistic pathogens in the gut community of *R. flavipes*. By conducting experiments over a span of 14 days, where termite workers are fed filter papers treated with different boric acid concentrations, we recorded their mortality and observed resulting alterations in their microbiomes after 7 days of boric acid treatment.

## Materials and Methods

### Insects

A colony of *R. flavipes* was collected from a forested area in Raleigh, NC, for use in experiments. Several hundred termites were maintained in plastic containers and provided with moist soil (Nature’s Care Organic & Natural Potting Mix with Water Conserve, Miracle-Gro, Marysville, OH), and pine shims (0.8 cm x 3.5 cm x 20 cm). The container was humidified by spraying water every 3 to 5 days. Termite colonies were maintained in an environmental rearing room at 26 °C, 50% RH, on a 12:12 LD cycle.

### Boric Acid Bioassays

To administer boric acid to termites, filter paper discs (10 mm Whatman No 1, Cytvia, Marlborough, MA, USA) were initially stapled to plastic microscope coverslips (22 x 22 mm). These papers were subsequently treated with either 4 µl of distilled water (control) or 4 µl of boric acid solutions at various concentrations (0.125%, 0.25%, 0.5%, 1%, and 4% boric acid). This range of solutions resulted in the corresponding amounts of 5, 10, 20, 40, and 160 µg of boric acid per paper, or 6.4, 12.7, 25.5, 50.9, and 203.7 µg of boric acid per cm^2^ of filter paper, respectively. Following treatment, the papers were air-dried for 24 h in a convection oven (BOF-102, Being Scientific, Ontario, CA, USA) maintained at 32 °C.

Prior to use in experiments, the filter papers were weighed using a precision balance (Explorer EX224, Ohaus, Wood Dale, IL, USA) to enable estimation of the amount of paper consumed by the termites. The average weight of filter papers was 6.8 mg ± 0.065 mg. Post-experiment, the papers were dried for 24 h at 32 °C, then weighed again. Their final weight was subtracted from the initial weight to determine the amount of paper consumed during the bioassay.

For experiments, worker termites were removed from the colony using an aspirator. Groups of 10 worker termites were then placed in plastic Petri dishes (100 x 10 mm) containing 15 ml of sand moistened with 5 ml of tap water. All termites were starved for 24 h prior to the bioassay. Termites were then offered one filter paper treated with water (control treatment) or with a specific concentration of boric acid (three replicates per treatment). Mortality checks were conducted every 24 h over 14 days. Termites were deemed to be dead if they did not exhibit any movement in response to gentle prodding with feather-tip forceps. Dead termites were removed from the Petri dish to prevent necrophagy, which could potentially confound the results pertaining to direct consumption of boric acid. Bioassays were conducted in incubators at 26°C, 90% RH, in continuous darkness. Termites fed untreated filter papers served as controls, and each treatment was replicated three times.

### Analysis of Termite Survivorship and Consumption of Boric Acid-Treated Papers

Mortality data for termites fed different concentrations of boric acid during the 14-day experimental period were analyzed using Kaplan-Meier survival analysis in SPSS 27 (IBM, Armonk, NY, USA). Different concentrations and controls were compared in a pair-wise manner using a log-rank test. Insects that survived beyond the 14-day period were right censored. To determine the hazard ratios for different treatments, a single proportional hazard regression was conducted. The water controls in each experiment were used as the baseline of comparison of hazard regression and statistical separation of different treatments. When possible, mean survival times (MSTs), median survival times, and relative log hazard ratios were estimated. The weight of treated filter papers was used to test for statistical differences between the amounts consumed in different treatments using Analysis of Variance (ANOVA) and a post-hoc t-tests (JMP 17, Cary, NC, USA).

### Gut Sampling, DNA Extraction and Sequencing

To investigate the impact of boric acid on the gut microbiome of *R. flavipes*, a separate set of termites from the same colony was utilized, following a setup similar to the previously described mortality assay. Termites were isolated using an aspirator and placed in Petri dishes. Twenty worker termites per Petri dish were starved for 24 h, and then provided two filter papers treated with either water or a specific concentration of boric acid, as described earlier. The use of 20 termites (compared to 10 in the mortality assays) and two filter papers ensured that sufficient live termites were available for microbiome sequencing. Mortality was monitored every 24 h for 7 days and any dead individuals were promptly removed to prevent necrophagy. Control treatments were filter papers treated with water and termites provided with no paper. Each treatment was replicated three times.

On the seventh day of the experiment, 10 worker termites from each replicate were anesthetized on ice packs for dissection. The gut was carefully removed by restraining the head with forceps and gently pulling the terminal segments of the abdomen with another forceps. Each termite gut was then placed in ZR BashingBead lysis tubes (Zymo Research, Irvine, CA, USA), suspended in 1 ml of ZR BashingBead buffer, and homogenized using a Benchmark Scientific D2400 Bead Beater (Sayreville, NJ, USA) for 4 cycles at 6.0 m/s for 11 seconds followed by 30 s of rest between each cycle. The homogenized gut samples were then stored at-80°C until DNA extraction. Each treatment and control group were replicated three times.

DNA from the homogenized termite guts was extracted using the Quick-DNA™ Fecal/Soil Microbe Microprep Kit (Zymo Research) following kit instructions. The bacterial community in the dissected guts of *R*. *flavipes* from control and boric-acid-treated groups was characterized via amplicon sequencing of the 16S rRNA gene. Polymerase Chain Reaction (PCR) Primers S-D-Bact-0341-b-S-17 (5′-CCTACGGGNGGCWGCAG-3′) and S-D-Bact-0785-a-A-21 (5′-GACTACHVGGGTATCTAATCC-3; Klindworth et al. 2013), each with a unique 8-bp-barcode at the 5’ end, were utilized to amplify the V3-V4 region of the 16S rRNA gene. PCR was performed with 50 μl reactions prepared with 10–50 ng of genomic DNA, 0.4 μM forward and reverse primers, 25 μl Taq polymerase master mix red (PCR Biosystems, Wayne, PA, USA), and 20 μl molecular-grade water. The following PCR program was used: 30 s of denaturation at 95°C, followed by 25 cycles of 20 s at 95°C, 20 s at 58°C, 30 s at 72°C, and a final elongation step at 72°C for 3 min. Amplicons were then cleaned using the DNA Clean and Concentrator-5 Kit (Zymo Research) following the manufacturer’s instructions. Purified amplicons were quantified and commercially sequenced at Novogene (Beijing, China) using the Illumina MiSeq platform.

### Microbiome Analysis

Processing and analysis of sequence data was conducted in line with the methodologies outlined by Schwarz et al. (2023), utilizing *mothur* (Schloss et al. 2009) for sequence processing and stringent quality control measures, followed by chimera removal and taxonomic classification. Briefly, we excluded contigs displaying ambiguities or homopolymer regions exceeding 10 bases. Additionally, sequences shorter than 200 bases, or those with an average quality score below 25, or a window average of 25 (over a window size of 50), were also eliminated. The remaining sequences were then aligned against the SILVA group’s comprehensive 50,000-position small subunit rRNA gene alignment. After the removal of chimeric sequences, the high-quality sequences were classified using the RDP Naïve Bayesian Classifier (Wang et al. 2004) implemented in *mothur*, referencing the SILVA database for taxonomic information (Quast et al. 2013). Following classification, sequences identified as originating from chloroplast or mitochondria were excluded from the subsequent ecological analyses.

To compare the alpha diversity of the termite microbiome across varying concentrations of boric acid, an ANOVA was conducted using the R (R Core Team 2021) package *vegan* (Dixon 2003) on the taxonomic composition of samples at the genus level. Beta diversity among the bacterial communities was assessed at the genus level using the Morisita-Horn metric via the vegan package. The resulting distance matrix was analyzed through two-dimensional non-metric multidimensional scaling (NMDS) (using the metaMDS function in *vegan*) to visualize microbial community dissimilarities. Additionally, permutational ANOVA (PERMANOVA) was performed on the Morisita-Horn distances to statistically evaluate the differences in termite gut microbiome compositions across varying concentrations of boric acid.

The random forest model, using the *randomForest* package (Liaw et al. 2002) in R, was employed to identify the top 30 bacterial genera contributing to the observed differences in the termite gut microbiome under varying boric acid treatments. The model, built using genus-level abundance data, ranked the taxa based on their importance measured by the Mean Decrease in Gini Index, ensuring the identification of the most influential genera. The log-transformed relative abundance of the top 30 taxa were then visualized using a heatmap (*pheatmap* package; Kolde and Kolde 2015), highlighting their distribution across different treatment groups and providing a clear representation of the taxa driving the observed differences in microbial community structure.

## Results

### Survival of *R. flavipes* After Feeding Upon Boric Acid-Treated Papers

Boric acid caused mortality in termites when they fed upon filter papers treated with different doses. Survivorship of *R. flavipes* was significantly impacted by the concentration of boric acid in a dose-dependent manner (Fig. 1, Table 1), with different treatments significantly impacting termite survivorship (Overall model, Chi-square = 48.1, d.f. = 5, *P* < 0.005). In comparisons between treatments, we found no significant differences between the control and 5 µg of boric acid per paper, the lowest tested dose (Chi-square = 1.9, d.f. = 1, *P* = 0.16). Termites being fed filter papers treated with 10, 15, or 20 µg of boric acid exhibited similar mortality rates (Chi-square = 0.001 ̶ 2.2, d.f. = 1, *P* > 0.05), but they were significantly different from the lower dose as well as from the highest dose. The highest mortality was observed in the 160 µg boric acid treatment, which was significantly different from all other treatments (Chi-square = 8.7 ̶ 29.5, d.f. = 1, *P* < 0.003). Some minimal mortality occurred after one day, but survivorship in the 160 µg boric acid treatment began to decline on day 4 and continued to decline until the end of the experiment. The median survival time in the 160 µg boric acid treatment was 7 days, which was therefore selected as the sampling point for the microbiome analysis in subsequent experiments. In the lower dose treatments, mortality began on days 6 or 7.

**Fig. 1.**
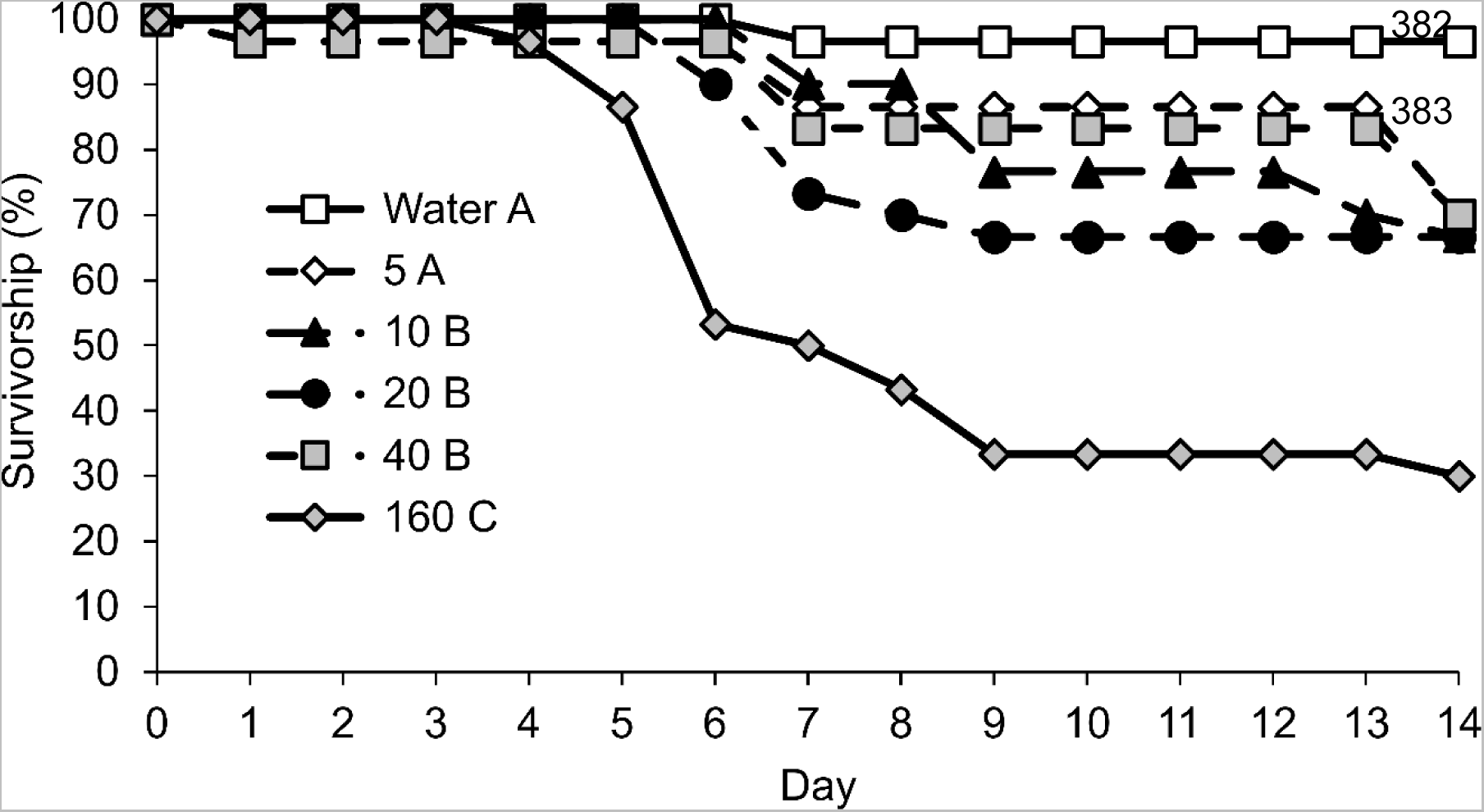
Mean proportional survival over time for *R. flavipes* individuals that were fed filter paper with water and varying amounts of boric acid (in µg). Each treatment represents 30 termites (3 replicates, each with 10 workers). Differences in the median survival time determined by a log-rank test are represented by the letters in the figure legend. Treatments that are not connected by the same letter are significantly different (*P* < 0.05).

**Table 1.** Kaplan-Meier estimates of mean survival time ± standard error for *R. flavipes* workers that were fed filter papers treated with different amounts of boric acid.

| Treatment | Grouping<sup>a</sup> | Mortality (%) | Mean Survival Time ±SE (Days) | Median Survival Time (Days) | Relative Log Hazard Ratio (95% CI)<sup>b</sup> |
| --- | --- | --- | --- | --- | --- |
| Control | A | 3.3 | 13.8 ±0.3 | $\geq 14$ | - |
| 5 μg | A | 13.3 | 12.8 ±0.6 | $\geq 14$ | 4.2 (0.5–38.1) |
| 10 μg | B | 30.0 | 12.4 ±0.5 | $\geq 14$ | 9.7 (1.2–76.5) |
| 20 μg | B | 33.3 | 11.7 ±0.6 | $\geq 14$ | 11.8 (1.5–92.0) |
| 40 μg | B | 30.0 | 12.7 ±0.6 | $\geq 14$ | 9.8 (1.2–77.2) |
| 160 μg | C | 70.0 | 9.1 ±0.7 | 7 | 34.2 (4.5–255) |

<sup>a</sup>Significantly different treatments as determined by a log-rank test are indicated by different letters in the grouping column.
<sup>b</sup>Relative log hazard ratios are not reported (denoted by a dash) for treatments used as the baseline for comparisons.

When termites were provided with filter papers treated with boric acid, the amount of paper consumed varied depending on the boric acid concentration present (Fig. 2, ANOVA, d.f.= 5, 17, *F* = 3.19, *P* = 0.046). Post-hoc *t*-tests revealed significant differences between treatments, with the control and the 5 µg treatment groups being statistically similar (*P* > 0.05), with all the paper being consumed in two replicates of the control and one replicate of the 5 µg treatment group. The 10 and 20 µg boric acid treatment groups had similar consumption to all other treatment groups, with an average consumption of 2.5 ± 2.0 and 3.7 ± 1.7 mg, respectively. Termites in the 40 and 160 µg treatment groups consumed the least amount of paper at an average 0.67 ± 0.3 and 0.77 ± 0.33 mg per treatment, respectively.

**Fig. 2.**
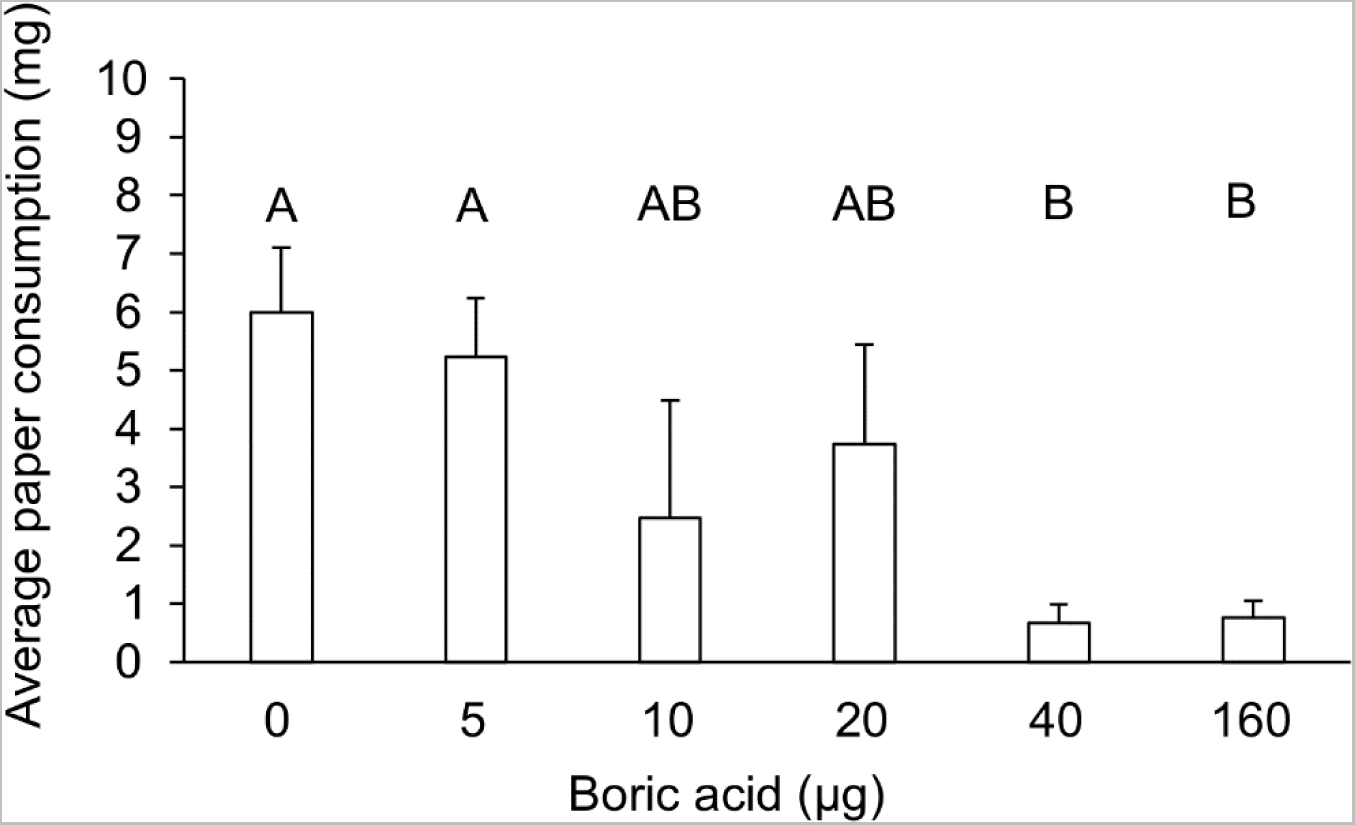
Average (±SEM) amount of paper consumed by *R. flavipes* based on treatment type. The 0 µg treatment served as the control group that was fed filter paper treated with water. Statistically significant differences in the average amount of paper consumed are indicated by different letters above the bars. Treatments not sharing the same letter are significantly different (ANOVA, *t*-test, *P* < 0.05).

### Termite Gut Bacterial Community in Relation to Boric Acid Concentration

We assessed the ability of boric acid to alter the microbial community of termites by feeding them different concentrations which also significantly impacted their survivorship. Data processing in *mothur* yielded a total of 9.87 million sequences from the 18 samples, with 548,456 sequences on average obtained per sample. 79% of all sequences (from all samples) were classified to the genus level using the Silva (v138) non-redundant database. This high classification success allowed us to conduct all downstream ecological analyses at the bacterial genus level. ANOVA comparing alpha diversity across boric acid concentrations revealed a significant (*P* < 0.005) difference supported by a high F-value (23.89). The PERMANOVA results indicate a highly significant effect of boric acid concentration on the termite gut microbiome structure (*F* = 46, R^2^ = 0.7417, *P* = 0.001), which suggest that around 74% of the observed variation in community structure can be explained by the grouping variable, i.e. boric acid concentration.

In the NMDS ordination plot (Fig. 3; Stress = 0.045) based on Morisita-Horn distances, a distinct separation is evident between the control group and the samples subjected to the highest boric acid dose (160 µg per filter paper). This separation underscores the significant disruption of bacterial community structure at this boric acid level, as further supported by the substantial proportion of explained variance in the PERMANOVA analysis. While there is a visible gradient across the boric acid doses, the absence of a distinct separation between the control group and the 40 µg boric acid treatment group suggests that lower concentrations of boric acid may only minimally affect the gut microbiome structure (as indicated by the overlapping points in the NMDS plot). This observation aligns with the PERMANOVA result, where the high R^2^ value indicates that higher concentrations of boric acid are driving the structural changes in the bacterial microbiome.

**Fig. 3.**
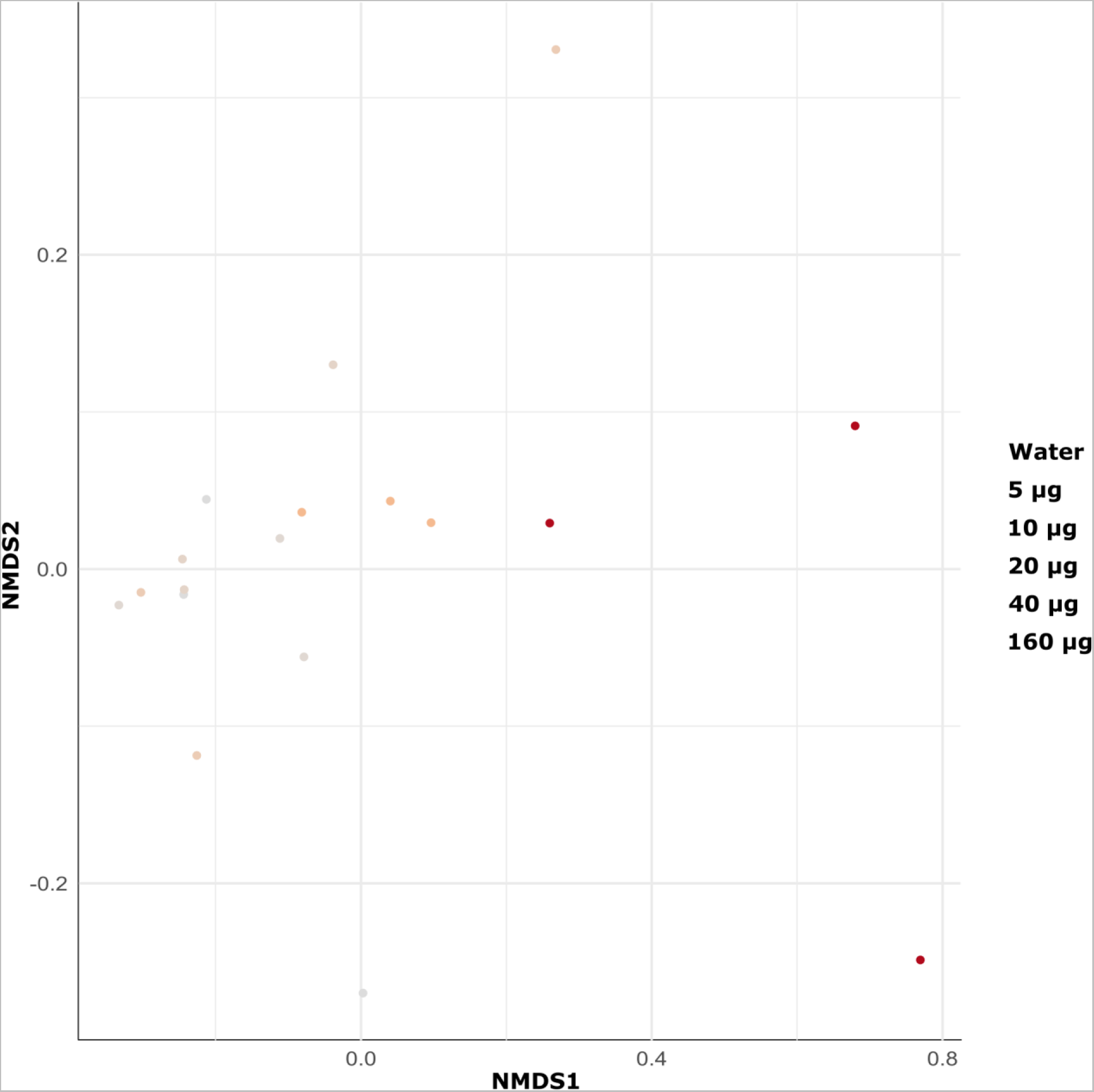
Non-metric multidimensional scaling (NMDS) plot depicting the structural dissimilarities (based on the Morisita-Horn metric) between termite gut bacterial communities in response to feeding upon filter papers treated with varying boric acid amounts. The two-dimensional solution had a low stress value (0.045), indicating a good representation of the data in reduced dimensions. Each point represents the compositional similarity of the termite gut communities, with closer points indicating more similar bacterial community compositions.

### Exposure to Boric Acid Causes Taxonomic Shifts in Termite Gut Community

To uncover specific taxonomic shifts in the termite microbiome in response to the boric acid treatments, we used a Random Forest machine learning approach. We selected the top 30 genera that exhibited marked shifts in relative abundance across the gradient of boric acid treatments (visualized in the heatmap in Fig. 4). The heatmap reveals several patterns in the distribution and abundance of bacterial taxa in relation to boric acid exposure in termites (for a more comprehensive exploration of the changes in the gut microbiome at various taxonomic levels, see Table S1). Notably, genera within the phylum Proteobacteria, including *Pseudomonas*, *Citrobacter*, and *Stenotrophomonas*, showed pronounced variations in relative abundance, particularly at the higher dose of 160 µg. Conversely, some members of the phylum Actinobacteriota, like the “Coriobacteriales *incertae sedis*” and the “uncultured *Raoultibacter*” group, displayed a decrease in relative abundance with increasing concentration of boric acid, with the lowest presence observed in the 160 µg treatment. Phyla such as the Fibrobacteraceae from the Fibrobacterota, which have been associated with cellulose degradation in the guts of higher termites, show a reduction in abundance with increasing boric acid.

**Fig. 4.**
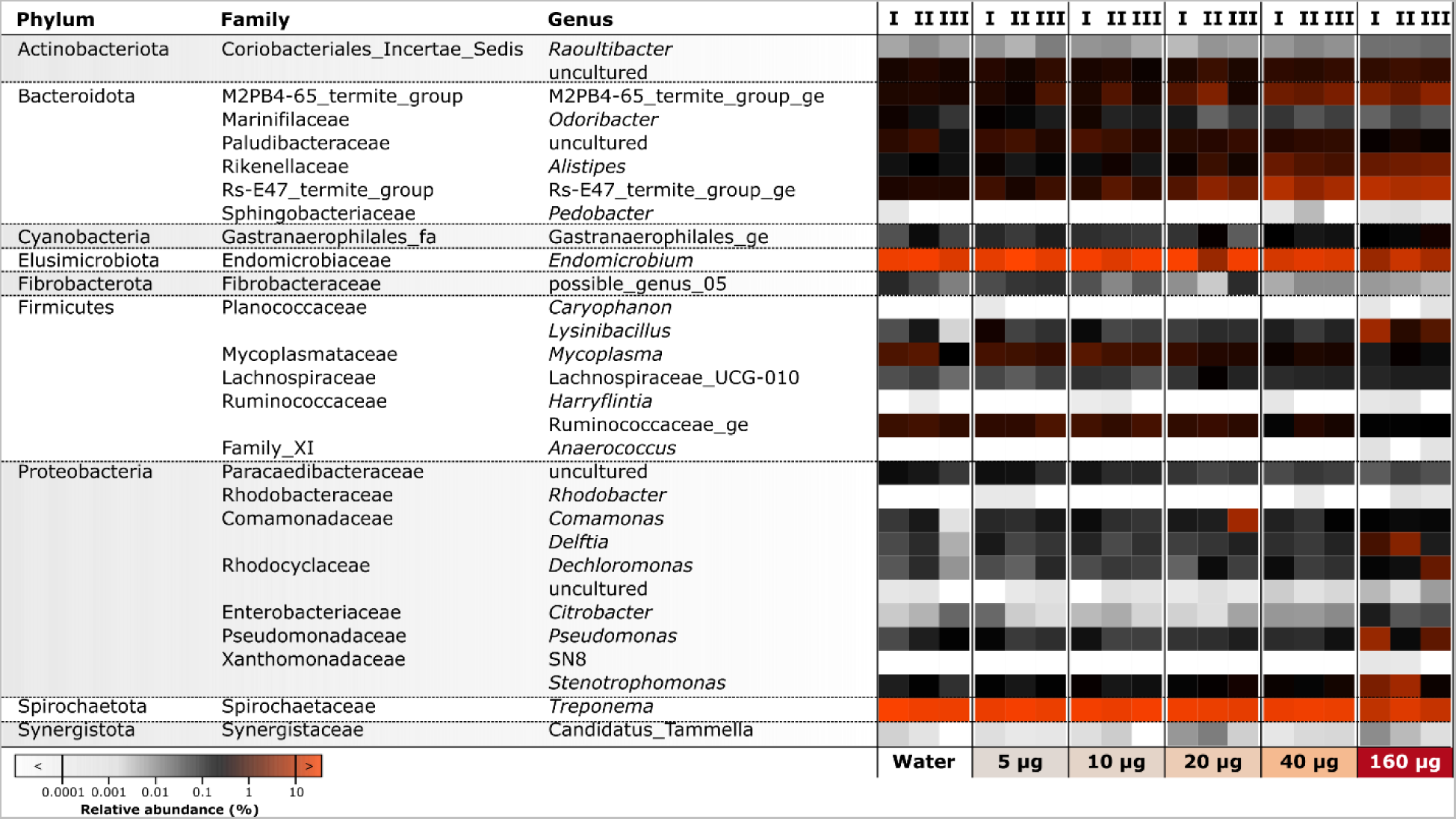
Heatmap depicting the distribution and abundance of the top 30 bacterial genera in the gut microbiome of *R. flavipes*, ranked by their contribution to group differentiation as determined by a random forest model. Each column represents a replicate of a treatment group (indicated below the heatmap), and variations in color intensity correspond to the log-transformed relative abundance of each genus, illustrating the impact of different concentrations of boric acid on bacterial community structure. For a more detailed exploration of changes in bacterial community structure, see Table S1.

Linear regression analysis further allowed us to interrogate the impact of increasing boric acid concentrations on the distribution and abundance of various bacterial taxa within the termite gut. It unveiled both significant positive and negative correlations across different bacterial lineages. Broad patterns in gut community structure reflecting boric acid concentrations were observed already at the phylum level. Bacteroidota and Firmicutes showed a strong positive correlation with increasing boric acid concentration, evidenced by their *P*-values (0.0102 and 0.0437, respectively) and R^2^ values (0.41 and 0.28, respectively). Conversely, Spirochaetota and Elusimicrobiota displayed significant negative trends, with *P*-values of 1.75e-05 and 0.0051, respectively. The high R^2^ values for Spirochaetota (0.77) and Elusimicrobiota (0.47) indicate that a significant portion of the variance in abundance for these phyla can be attributed to boric acid concentrations.

Diving deeper at the genus level, *Alistipes* (Phylum: Bacteroidota) demonstrated a significant positive correlation with boric acid concentration (*P* = 1.47e-05), with an R^2^ value of 0.78. The abundance of *Alistipes* increased from 0.46% ± 0.03% in water-treated samples to 2.97% ± 1.05% in samples treated with 160 µg of boric acid. *Raoultibacter* (Phylum: Actinomycetota), despite its significant shift (*P* = 3.23e-05, R^2^ = 0.75), remained a comparatively rare taxon, with a relative abundance of less than 0.05% across all treatments.

Significant positive trends were also identified for several other taxa, including “Rs-E47_termite_group_ge” (Phylum: Bacteroidota), *Lysinibacillus* and *Mycoplasma* (Phylum: Firmicutes), as well as *Pseudomonas*, *Delftia*, *Dechloromonas*, and *Stenotrophomonas* (Phylum: Proteobacteria). These taxa demonstrated substantial increases in mean relative abundance— ranging from 18-fold to 45-fold—between control filter papers and those supplemented with 160 µg of boric acid. For instance, *Lysinibacillus* surged from 0.06% ± 0.07% in the control to 2.53% ± 2.76% in termites exposed to boric acid. Similarly, the abundance of Rs-E47_termite_group_ge escalated from 0.48% ± 0.04% to 8.43% ± 1.54%.

Only one of the genera shortlisted by our random forest approach, *Endomicrobium,* presented a significant negative trend (*P* = 0.00512, R^2^ = 0.47), indicating a drop in abundance with rising boric acid concentrations. Its relative abundance diminished from 26.38% ± 6.36% in the control group to 8.32% ± 4.39% in the 160 µg boric acid treatment group, highlighting a considerable impact of boric acid on this taxon’s presence within the termite gut.

## Discussion

Our study builds upon existing evidence to further validate boric acid’s lethal impact on termites, with survivorship in the eastern subterranean termite *R. flavipes* dropping in a dose-dependent manner (Fig. 1, Table 1). This finding supports previous observations of dose-dependence in *R. flavipes* (Su et al. 1994), and the termites *C. formosanus* (Su et al. 1994; Gentz et al. 2009; Gentz and Grace 2009) and *Heterotermes indicola* (Wasmann) (Blattodea: Rhinotermitidae) (Farid et al. 2015). The highest amount tested in our study was 160 µg of boric acid per 6.8 mg of filter paper, which corresponds to approximately 23,529 ppm (calculated as 160 µg/6800 µg to convert to ppm). This concentration led to significant termite mortality, which parallels the mortality observed at lower concentrations (10,000 ppm) reported by Farid et al. (2015). This dose-dependent effect has been similarly documented for boric acid when ingested by *Blattella germanica* L. (Blattodea: Ectobiidae) (Cochran 1995; Habes et al. 2001; Gore and Schal 2004; Habes et al. 2006; Jiang et al. 2021), *Galleria mellonella* L. (Lepidoptera: Pyralidae) (Gwokyalya and Altuntaş 2019), *Apis mellifera* L. (Hymenoptera: Apidae) (da Silva Cruz et al. 2010), and *Cimex lectularius* L. (Hemiptera: Cimicidae) (Sierras et al. 2018).

We found that termites consumed less paper in a dose-dependent manner, with the least amount consumed from papers treated with the highest concentrations of boric acid (Fig. 2). Although boric acid does not cause an immediate repellent effect, our results and those of Farid et al. (2015) show that it ultimately leads to reduced feeding. Other studies have shown that *R. flavipes*, *C. formosanus* and *Coptotermes gestroi* (Wasmann) (Blattodea: Rhinotermitidae), consume less wood when treated with higher concentrations boric acid solutions (Casarin et al. 2009, Kard 2001). While one might speculate that reduced feeding was due to termite mortality, this scenario is unlikely as the 40 µg treatment had lower mortality than the 160 µg treatment, despite similar amounts of filter paper consumption.

While the specific mode(s) of action of boric acid in termites remains underexplored, evidence from prior studies with termites and several other insect species indicates its significant impact on the digestive system. Studies, including Cochran (1995) and Habes et al. (2005), have demonstrated that boric acid induces structural changes in the midgut of *B. germanica*, potentially impairing nutrient absorption. It was later shown by Yang et al. (2021) that boric acid can create pores in the midgut lining of *B. germanica*, facilitating infections by *Metarhizium anisopliae* (Metchnikoff) (Hypocreales: Clavicipitaceae) spores. Given that cockroaches and termites are close phylogenetic relatives, it is plausible that termites experience similar digestive disruptions. However, the potential impairment of the midgut has also been observed in Argentine ant *Linepithema humile* (Mayr; Hymenoptera: Formicidae) (Klotz et al. 2002) and the honey bee *A. mellifera* (da Silva Cruz et al. 2010), underscoring the disruption of digestive physiology as an significant contributor to the insecticidal effects of boric acid.

Our results reveal that boric acid induces significant alterations in alpha diversity and structural changes within the termite gut microbiome (Fig. 3). This aligns with observations by Jiang et al. (2021), where boric acid was shown to disrupt the normal gut microbiome composition in *B. germanica*, suggesting a similar mechanism might be at play in termites. The decrease in alpha diversity and significant shifts in microbial community composition, especially at higher boric acid concentrations, are indicative of dysbiosis, defined as an imbalance or shift in a naturally present microbiome of a healthy host (Petersen and Round 2014). Given the reliance of termites on their gut microbiome for the symbiotic digestion of wood, such an imbalance seems to magnify the detrimental effects of boric acid.

A loss of diversity in the gut microbiome is typically tied to reduced functionality, and is often accompanied by a loss of beneficial symbionts and an expansion of opportunistic pathogens (Petersen and Round 2014). The clear clustering and separation observed in the NMDS plot for termites exposed to filter paper treated with 160 µg of boric acid compared to the control group illustrates the significant impact of boric acid on gut community structure (Fig. 3). This pattern not only confirms the presence of dysbiosis but also suggests that higher concentrations of boric acid may lead to more pronounced shifts in microbial populations, potentially disrupting critical processes such as digestion, nutrient absorption, and pathogen resistance within the termite gut. However, the noticeable dispersal in clustering among replicates treated with higher concentrations of boric acid (Fig. 3) indicates variability in the response of the termite gut microbiome to the treatment. This variability, coupled with the observed shifts in the relative abundance of certain bacterial genera (Fig. 4), suggests that the dysbiotic effects of boric acid on the termite gut microbiome are not deterministic and may vary depending on a complex range of factors.

Our screening for microbial taxa contributing to the overall differences observed in microbiome structure revealed a downward trend (with increasing boric acid exposure) in the abundance of key genera, such as *Endomicrobium* (Fig. 4) that are major endosymbionts of *Trichonympha* flagellates. The drop in the relative abundance of these flagellate endosymbionts reinforces the findings of Kard (2001), who found boric acid at 4% solution (equivalent to our 160 µg treatment) to be effective at disrupting the flagellate community of *R. flavipes*. Other forms of defaunation (experimental removal of flagellates) by force-feeding starch or other treatments have also been shown to markedly reduce the abundance of this bacterial endosymbiont (Ikeda-Ohtsubo et al. 2010; Mikaelyan et al. 2017). The observed negative correlation between boric acid concentration and the Spirochaetota phylum, essential for providing vital microbial services (Tokuda 2021), such as lignocellulose degradation (Mikaelyan et al. 2014; Tokuda et al. 2018), nitrogen fixation (Lilburn et al. 2001; Ohkuma et al. 2015), and homoacetogenesis (Leadbetter et al. 1999; Ottesen and Leadbetter 2011) highlights that its depletion poses a significant detriment to termite metabolism.

At the same time, the observed increase in certain genera, notably *Lysinibacillus*, *Pseudomonas*, and *Mycoplasma*, suggests a proliferation of opportunistic pathogens. *Lysinibacillus sphaericus* (previously known as *Bacillus sphaericus*) is recognized for its mosquito larvicidal properties (Chalegre et al. 2015; Rezende et al. 2019), and is closely related to several insect-pathogenic species (Nakamura 2000). The genus *Pseudomonas* similarly includes many pathogens of arthropods (Roobakkumar et al. 2011; Carrau et al. 2021; Hamze et al. 2022) with preadaptations to infect insects as specialists or as opportunists. When *Mastotermes darwiniensis* Froggatt (Blattodea: Mastotermitidae) is in a state of dysbiosis due to starch feeding, there is an increase in bacteria in their hindguts, that may be opportunists (Veivers et al. 1983). In *B. germanica*, boric acid exposure led to dysbiosis, characterized by a decrease in beneficial bacteria like *Bacteroides* and *Enterococcus*, and an uptick in potentially pathogenic *Weissella* species (Jiang et al. 2021). Similarly, when *Drosophila melanogaster* Meigen (Diptera: Drosophilidae) are in a state of dysbiosis, opportunistic bacteria in their guts, such as *Gluconobacter mortifer*, become pathogens (Lee and Lee 2014).

The dysbiosis we observed could potentiate the vulnerability of termites to external entomopathogens as well, as demonstrated by Yang et al. (2021), where boric acid enhanced the virulence of *M. anisopliae* against *B. germanica* by altering the gut microbiome. This suggests that boric acid’s mode of action may extend to facilitating the invasion and proliferation of pathogenic microbes within the termite gut, offering a novel perspective on utilizing boric acid in integrated pest management (IPM) strategies that include baiting tactics.

The decrease in alpha diversity and significant shifts in microbial community composition, especially at higher concentrations of boric acid, indicate a disruption of the gut microbiome’s equilibrium. This dysbiosis could further compromise termite health and resilience, beyond the direct toxicological effects of boric acid. Considering the substantial alterations in the gut microbiome and the increased termite mortality associated with boric acid consumption, our findings support the exploration of synergistic approaches for termite management. Our study contributes to the growing body of evidence on the utility of boric acid in termite management, not only as a direct toxicant but also as a disruptor of gut microbiome homeostasis. Future research should focus on elucidating the specific mechanisms by which boric acid induces dysbiosis and exploring its synergistic potential with microbial control agents, paving the way for innovative and sustainable termite control strategies.

## Author Contributions

Conceptualization, methodology, software, validation, writing—original draft preparation, and writing—review and editing, A.R.A., M.S., C.S., and A.M.; investigation, formal analysis, and data curation, A.R.A., M.S., and A.M; resources, supervision, project administration, and funding acquisition A.M. and C.S.; visualization, A.R.A., M.S., and A.M.

## Data Accessibility

The 16S rRNA gene sequence reads analyzed as part of this study are available in the NCBI Sequence Read Archive (SRA) repository under Bioproject ID PRJNA1096953.

## Supplementary data

Table S1. An expandable Excel spreadsheet for an in-depth analysis of the distribution and relative abundance of bacterial taxa in the gut microbiome of *Reticulitermes flavipes*, in response to boric acid ingestion.

## References

1. Brune A. Symbiotic digestion of lignocellulose in termite guts. Nat. Rev. Microbiol. 2014:12(3):168–180. 10.1038/nrmicro3182.

2. Brune A, Dietrich C. The gut microbiota of termites: digesting the diversity in the light of ecology and evolution. Annu. Rev. Microbiol. 2015:69:145–166. 10.1146/annurev-micro-092412-155715.

3. Casarin FE, Costa-Leonardo AM, Bueno OC. Laboratory assessment of two active ingredients for control of *Coptotermes gestroi* (Isoptera: Rhinotermitidae). Sociobiol. 2009:54(3), 787–789.

4. Carrau T, Thümecke S, Silva LM, Perez-Bravo D, Gärtner U, Taubert A, Hermosilla C, Vilcinskas A, Lee KZ. The Cellular innate immune response of the invasive pest insect *Drosophila suzukii* against *Pseudomonas entomophila* involves the release of extracellular traps. Cells. 2021:10(12):3320. 10.3390/cells10123320.

5. de Melo Chalegre KD, Tavares DA, Romão TP, de Menezes HS, Nascimento NA, de Oliveira CM, de-Melo-Neto OP, Silva-Filha MH Co-selection and replacement of resistance alleles to *Lysinibacillus sphaericus* in a *Culex quinquefasciatus* colony. FEBS J. 2015:282(18):3592– 3602. 10.1111/febs.13364

6. Cochran DG. Toxic effects of boric acid on the German cockroach. Experientia. 1995:51:561–3. 10.1007/BF02128743

7. R Core Team. R: A language and environment for statistical computing. Vienna: R foundation for statistical computing. R Foundation for Statistical Computing. 2021. Available from https://cir.nii.ac.jp/crid/1370576118723163397.

8. Dedeine F, Dupont S, Guyot S, Matsuura K, Wang C, Habibpour B, Bagnères AG, Mantovani B, Luchetti A. Historical biogeography of *Reticulitermes termites* (Isoptera: Rhinotermitidae) inferred from analyses of mitochondrial and nuclear loci. Molecular Phylogenetics and Evolution. 2016:94:778–90. 10.1016/j.ympev.2015.10.020

9. Dixon P. VEGAN, a package of R functions for community ecology. J. Veg. Sci. 2003:14(6):927–930.

10. Ebeling W. Urban Entomology, Division of Agricultural Sciences. University of California, Berkeley; 1978.

11. Farid A, Zaman M, Saeed M, Khan M, Shah TB. 2015. Evaluation of boric acid as a slow-acting toxicant against subterranean termites (*Heterotermes* and *Odontotermes*). Journal of entomology and zoology studies. 2015:3:213–216.

12. Gentz MC, Grace JK. A review of boron toxicity in insects with an emphasis on termites. J. Agric. Urban Entomol. 2006:23(4):201–207.

13. Gentz MC, Grace JK. The response and recovery of the Formosan subterranean termite (Coptotermes formosanus Shiraki) from sublethal boron exposures. Int. J. Pest Manage. 2009:55(1):63–67. 10.1080/09670870802460721.

14. Gentz MC, Grace JK, Mankowski ME. Horizontal transfer of boron by the Formosan subterranean termite (Coptotermes formosanus Shiraki) after feeding on treated wood. Holzforschung. 2009:63(1):113–117. 10.1515/HF.2009.025.

15. Gile GH. Protist symbionts of termites: diversity, distribution, and coevolution. Biol. Rev. Camb. Philos. Soc. 2024:99(2):622–652. 10.1111/brv.13038.

16. Gore JC, Schal C. Laboratory evaluation of boric acid-sugar solutions as baits for management of German cockroach infestations. J. Econ. Entomol. 2004:97(2):581–587 10.1093/jee/97.2.581.

17. Gwokyalya R, Altuntaş H. Boric acid-induced immunotoxicity and genotoxicity in model insect *Galleria mellonella* L. (Lepidoptera: Pyralidae). Arch. Insect Biochem. Physiol. 2019:101(4):e21588.

18. Habes D, Kilani-Morakchi S, Aribi N, Farine JP, Soltani N. Toxicity of boric acid to *Blattella germanica* (Dictyoptera: Blattellidae) and analysis of residues in several organs. Meded. Rijksuniv. Gent Fak. Landbouwkd. Toegep. Biol. Wet. 2001:66(2a):525–534.

19. Habes D, Morakchi S, Aribi N, Farine JP, Soltani N. Boric acid toxicity to the German cockroach, *Blattella germanica*: Alterations in midgut structure, and acetylcholinesterase and glutathione S-transferase activity. Pestic. Biochem. Physiol. 2006:84(1):17–24. 10.1016/j.pestbp.2005.05.002

20. Hamze R, Nuvoli MT, Pirino C, Ruiu L. Compatibility of the bacterial entomopathogen *Pseudomonas protegens* with the natural predator *Chrysoperla carnea* (Neuroptera: Chrysopidae). J. Invertebr. Pathol. 2022:194:107828. 10.1016/j.jip.2022.107828

21. Ikeda-Ohtsubo W, Faivre N, Brune A. 2010. Putatively free-living “Endomicrobia”-ancestors of the intracellular symbionts of termite gut flagellates? Environ. Microbiol. Rep. 2(4):554–559. 10.1111/j.1758-2229.2009.00124.x

22. Jiang M, Dong F-Y, Pan X-Y, Zhang Y-N, Zhang F. 2021. Boric acid was orally toxic to different instars of Blattella germanica (L.) (Blattodea: Blattellidae) and caused dysbiosis of the, gut microbiota. Pestic. Biochem. Physiol. 172:104756. 10.1016/j.pestbp.2020.104756

23. Kard BM. 2001. Detrimental Effects of Boric-Acid-Treated Soil Against Foraging Subterranean Termites (Isoptera: Rhinotermitidae). Sociobiology. 2001:37(2):363-378.

24. Klindworth A, Pruesse E, Schweer T, Peplies J, Quast C, Horn M, Glöckner FO. Evaluation of general 16S ribosomal RNA gene PCR primers for classical and next-generation sequencing-based diversity studies. Nucleic Acids Res. 2013:41(1):e1–e1. 10.1093/nar/gks808.

25. Klotz JH, Amrhein C, McDaniel S. Assimilation and toxicity of boron in the Argentine ant (Hymenoptera: Formicidae). J. Entomol. Sci. 2002:37(2):193–199. 10.18474/0749-8004-37.2.193

26. Kolde R, Kolde MR. 2015. Package “pheatmap.” R package. 1(7):790.

27. Leadbetter JR, Schmidt TM, Graber JR, Breznak JA. Acetogenesis from H_2_ plus CO_2_ by spirochetes from termite guts. Science. 1999:283(5402):686–689. 10.1126/science.283.5402.686

28. Lee, K.-A., Lee, W.-J. *Drosophila* as a model for intestinal dysbiosis and chronic inflammatory diseases. Developmental & Comparative Immunology. 2014:42: 102 ̶ 110. 10.1016/j.dci.2013.05.005

29. Liaw A, Wiener M, Others. 2002. Classification and regression by randomForest. R news. 2(3):18–22.

30. Lilburn TG, Kim KS, Ostrom NE, Byzek KR, Leadbetter JR, Breznak JA. Nitrogen fixation by symbiotic and free-living spirochetes. Science. 2001:292(5526):2495–2498. 10.1126/science.1060281

31. Mikaelyan A, Strassert JFH, Tokuda G, Brune A. The fibre-associated cellulolytic bacterial community in the hindgut of wood-feeding higher termites (*Nasutitermes* spp.). Environ. Microbiol. 2014:16(9):2711–2722. 10.1111/1462-2920.12425

32. Mikaelyan A, Thompson CL, Meuser K, Zheng H, Rani P, Plarre R, Brune A. High-resolution phylogenetic analysis of Endomicrobia reveals multiple acquisitions of endosymbiotic lineages by termite gut flagellates. Environ. Microbiol. Rep. 2017:9(5):477–483. 10.1111/1758-2229.12565

33. Nakamura LK. Phylogeny of *Bacillus sphaericus*-like organisms. Int. J. Syst. Evol. Microbiol. 2000:50(5):1715–1722. 10.1099/00207713-50-5-1715

34. Ohkuma M, Noda S, Hattori S, Iida T, Yuki M, Starns D, Inoue JI, Darby AC, Hongoh Y. Acetogenesis from H_2_ plus CO_2_ and nitrogen fixation by an endosymbiotic spirochete of a termite-gut cellulolytic protist. Proc. Natl. Acad. Sci. 2015:112(33):10224–10230. 10.1073/pnas.1423979112

35. Oi F. 2022. A Review of the Evolution of Termite Control: A Continuum of Alternatives to Termiticides in the United States with Emphasis on Efficacy Testing Requirements for Product Registration. Insects. 13(1). 10.3390/insects13010050

36. Ottesen EA, Leadbetter JR. Formyltetrahydrofolate synthetase gene diversity in the guts of higher termites with different diets and lifestyles. Appl. Environ. Microbiol. 2011:77(10):3461–3467. 10.1128/AEM.02657-10

37. Petersen C, Round JL. Defining dysbiosis and its influence on host immunity and disease. Cell. Microbiol. 2014:16(7):1024–1033. 10.1111/cmi.12308

38. Protasov E, Nonoh JO, Kästle Silva JM, Mies US, Hervé V, Dietrich C, Lang K, Mikulski L, Platt K, Poehlein A, Köhler-Ramm T, Miambi E, Boga HI, Feldewert C, Ngugi DK, Plarre R, Sillam-Dussès D, Šobotník J, Daniel R and Brune A. Diversity and taxonomic revision of methanogens and other archaea in the intestinal tract of terrestrial arthropods. Front. Microbiol. 2023:14:1281628. 10.3389/fmicb.2023.1281628

39. Quast C, Pruesse E, Yilmaz P, Gerken J, Schweer T, Yarza P, Peplies J, Glöckner FO. The SILVA ribosomal RNA gene database project: improved data processing and web-based tools. Nucleic Acids Res. 2013:41(D1):D590–596. 10.1093/nar/gks1219

40. Rezende TM, Rezende AM, Luz Wallau G, Santos Vasconcelos CR, de-Melo-Neto OP, Silva-Filha MH, Romão TP. 2019. A differential transcriptional profile by Culex quinquefasciatus larvae resistant to *Lysinibacillus sphaericus* IAB59 highlights genes and pathways associated with the resistance phenotype. Parasit. Vectors. 2019:12(1):1–17. 10.1186/s13071-019-3661-y

41. Roobakkumar A, Babu A, Kumar DV, Sarkar S. 2011. Pseudomonas fluorescens as an efficient entomopathogen against Oligonychus coffeae Nietner (Acari: Tetranychidae) infesting tea. J Entomol Nematol. 2011:3(5):73-77

42. Rosengaus RB, Zecher CN, Schultheis KF, Brucker RM, Bordenstein SR. Disruption of the termite gut microbiota and its prolonged consequences for fitness. Appl. Environ. Microbiol. 2011:77(13):4303–4312. 10.1128/AEM.01886-10.

43. Rust MK, Su N-Y. Managing social insects of urban importance. Annu. Rev. Entomol. 2012:57:355–375. 10.1146/annurev-ento-120710-100634

44. Scharf ME. Challenges and physiological implications of wood feeding in termites. Curr Opin Insect Sci. 2020:41:79–85. 10.1016/j.cois.2020.07.007

45. Schloss PD, Westcott SL, Ryabin T, Hall JR, Hartmann M, Hollister EB, Lesniewski RA, Oakley BB, Parks DH, Robinson CJ, Sahl JW, Stres B, Thallinger GG, Van Horn DJ, Weber CF. Introducing mothur: open-source, platform-independent, community-supported software for describing and comparing microbial communities. Appl. Environ. Microbiol. 2009:75(23):7537–7541. 10.1128/AEM.01541-09.

46. Schubert DM. Boric Oxide, Boric Acid, and Borates. In: Ullmann’s Encyclopedia of Industrial Chemistry. Weinheim, Germany: Wiley-VCH Verlag GmbH & Co. KGaA. 2000:1–32. 10.1002/14356007.a04_263.pub2

47. Schwarz M, Beza-Beza CF, Mikaelyan A. Wood fibers are a crucial microhabitat for cellulose-and xylan-degrading bacteria in the hindgut of the wood-feeding beetle Odontotaenius disjunctus. Front. Microbiol. 2023:14. 10.3389/fmicb.2023.1173696

48. Sierras A, Wada-Katsumata A, Schal C. Effectiveness of boric acid by ingestion, but not by contact, against the common bed bug (Hemiptera: Cimicidae). J. Econ. Entomol. 2018:111(6):2772–2781. 10.1093/jee/toy260

49. da Silva Cruz A, da Silva-Zacarin EC, Bueno OC, Malaspina O. Morphological alterations induced by boric acid and fipronil in the midgut of worker honeybee (*Apis mellifera* L.) larvae: morphological alterations in the midgut of *A. mellifera*. Cell Biol. Toxicol. 2010:26:165–176. 10.1007/s10565-009-9126-x

50. Su N-Y, Tokoro M, Scheffrahn RH. Estimating Oral Toxicity of Slow-Acting Toxicants Against Subterranean Termites (Isoptera: Rhinotermitidae). J. Econ. Entomol. 1994:87(2):398–401.

51. Tokuda G. 2021. Origin of symbiotic gut spirochetes as key players in the nutrition of termites. Environmental microbiology. 23(8):4092–4097. 10.1111/1462-2920.15625

52. Tokuda G, Mikaelyan A, Fukui C, Matsuura Y, Watanabe H, Fujishima M, Brune A. Fiber-associated spirochetes are major agents of hemicellulose degradation in the hindgut of wood-feeding higher termites. Proc. Natl. Acad. Sci. 2018:115(51):E11996–E12004. 10.1073/pnas.1810550115

53. Veivers, P.C., O’Brien, R.W., Slaytor, M. Selective defaunation of *Mastotermes darwiniensis* and its effect on cellulose and starch metabolism. Insect Biochemistry. 1983:13:95 ̶101.

54. Wang Q, Garrity GM, Tiedje JM, Cole JR. Naive Bayesian classifier for rapid assignment of rRNA sequences into the new bacterial taxonomy. Applied and environmental microbiology. 2007:73(16):5261–5267. 10.1128/aem.00062-07

55. Watanabe H, Tokuda G. Cellulolytic systems in insects. Annu. Rev. Entomol. 2010:55:609–632. 10.1146/annurev-ento-112408-085319

56. Yang R, Zhang M, Schal C, Jiang M, Cai T, Zhang F. Boric acid enhances Metarhizium anisopliae virulence in Blattella germanica (L.) by disrupting the gut and altering its microbial community. Biol. Control. 2021:152:104430. 10.1016/j.biocontrol.2020.104430

